## Supplementary material for "Concurrent remodelling of nucleolar 60S subunit precursors by the Rea1 ATPase and Spb4 RNA helicase": Figure Supplements

### Figure Supplements & Supplementary Tables

Mitterer, Thoms *et al*

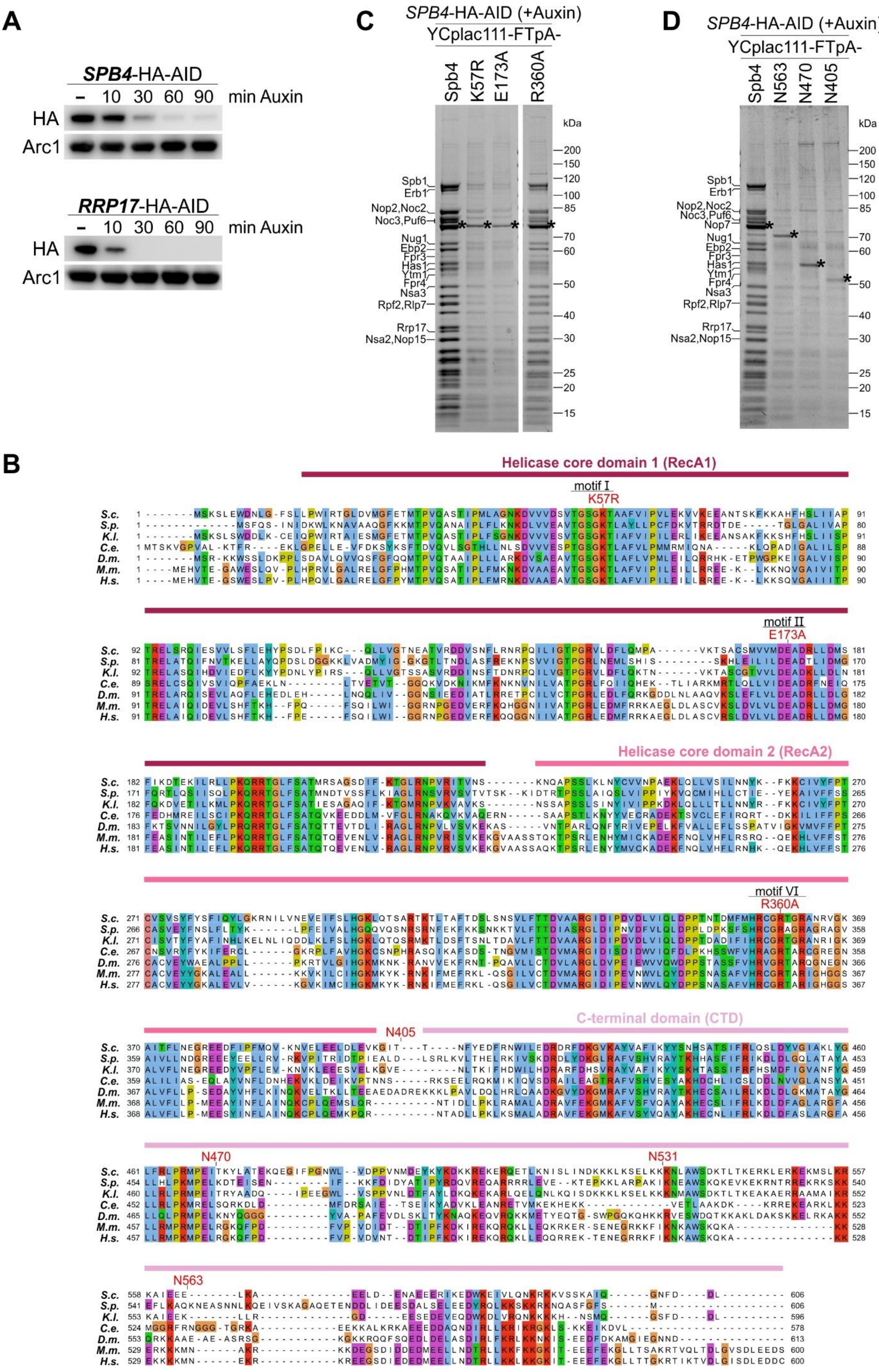
A
SPB4-HA-AID
– 10 30 60 90 min Auxin
HA
Arc1
RRP17-HA-AID
– 10 30 60 90 min Auxin
HA
Arc1
C
SPB4-HA-AID (+Auxin)
YCplac111-FTpA-
Spb4 K57R E173A R360A
kDa
200 150 120 100 85 70 60 50 40 30 25 20 15
Spb1
Erb1
Nop2,Noc2
Noc3,Puf6
Nug1
Ebp2
Fpr3
Has1
Ytm1
Fpr4
Nsa3
Rpf2,Rlp7
Rrp17
Nsa2,Nop15
D
SPB4-HA-AID (+Auxin)
YCplac111-FTpA-
Spb4 N563 N470 N405
kDa
Spb1
Erb1
Nop2,Noc2
Noc3,Puf6
Nop7
Nug1
Ebp2
Fpr3
Has1
Ytm1
Fpr4
Nsa3
Rpf2,Rlp7
Rrp17
Nsa2,Nop15
B
Helicase core domain 1 (RecA1)
motif I
K57R
motif II
E173A
Helicase core domain 2 (RecA2)
motif VI
R360A
C-terminal domain (CTD)
N405
N470
N531
N563
S.c.
S.p.
K.l.
C.e.
D.m.
M.m.
H.s.

**Figure 1–Figure Supplement 1.** (A) Spb4-HA-AID and Rrp17-HA-AID are efficiently degraded in the presence of auxin. Logarithmically growing *SPB4*-HA-AID and *RRP17*-HA-AID cells were treated with 0.5 mM auxin and the level of Spb4-HA-AID and Rrp17-HA-AID protein degradation was assessed over 90 min by western blot analysis of whole cell lysates using an anti-HA antibody. Equal sample loading was controlled by western blot analysis using an anti-Arc1 antibody. (B) Multiple sequence alignment of Spb4 orthologues. The Spb4 protein sequences from *Saccharomyces cerevisiae* (*S.c.*), *Schizosaccharomyces pombe* (*S.p.*), *Kluyveromyces lactis* (*K.l.*), *Caenorhabditis elegans* (*C.e.*), *Drosophila melanogaster* (*D.m.*), *Mus musculus* (*M.m.*), and *Homo sapiens* (*H.s.*) were aligned with Clustal omega and illustrated with Jalview. Point mutations introduced in catalytic helicase motifs and C-terminal truncation mutants are indicated. (C-D) Efficient pre-60S affinity purification via Spb4 is impaired in catalytic helicase mutants and C-terminal truncation mutants. Indicated plasmid-borne wild-type and mutant Spb4 variants fused to a C-terminal FTpA tag were affinity purified from cells upon auxin-induced depletion of chromosomal *SPB4*-HA-AID. Final eluates were analysed by SDS-PAGE and Coomassie staining.

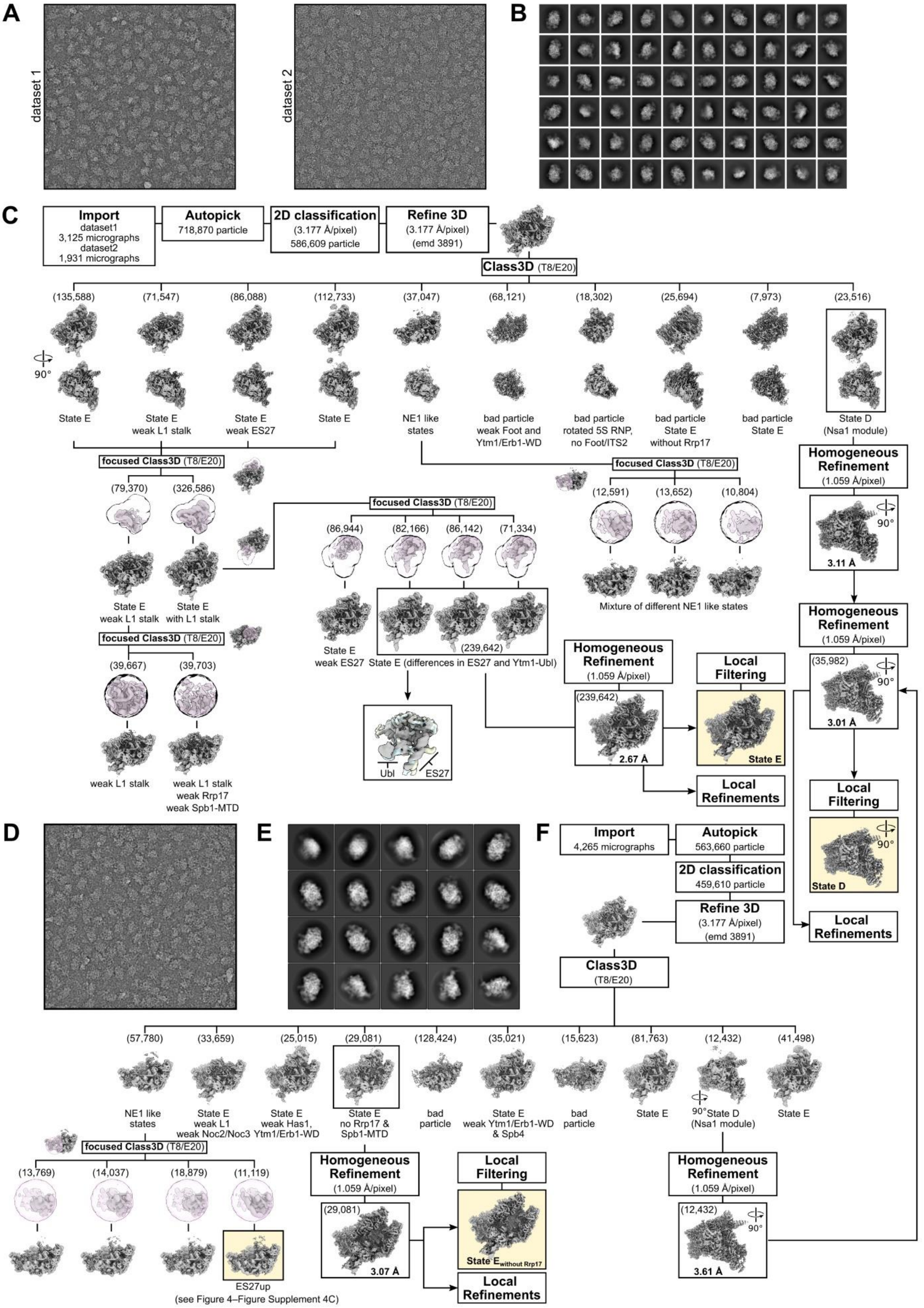


**Figure3–Figure Supplement 1. Cryo-electron microscopy data processing scheme.** (A-B) Representative micrographs and 2D class averages of the two datasets used for processing

of the Spb4-FTpA sample and determination of the state E structure. (C) Sorting scheme of the Spb4 sample. Final classes are indicated. (D-F) Representative micrograph (D), 2D classes (E) and processing scheme (F) of the sample used for determination of the state E-like structure without Rrp17/Spb1-MTD. Particles for state D from (C) and (F) were combined to increase the resolution of the final state D map. Particles of the NE1-like state with the ES27 rRNA in an upward conformation (F) were combined with an identical class described in Figure4–Figure Supplement 4C. Final cryo-EM volumes are highlighted and shown with their respective resolution.

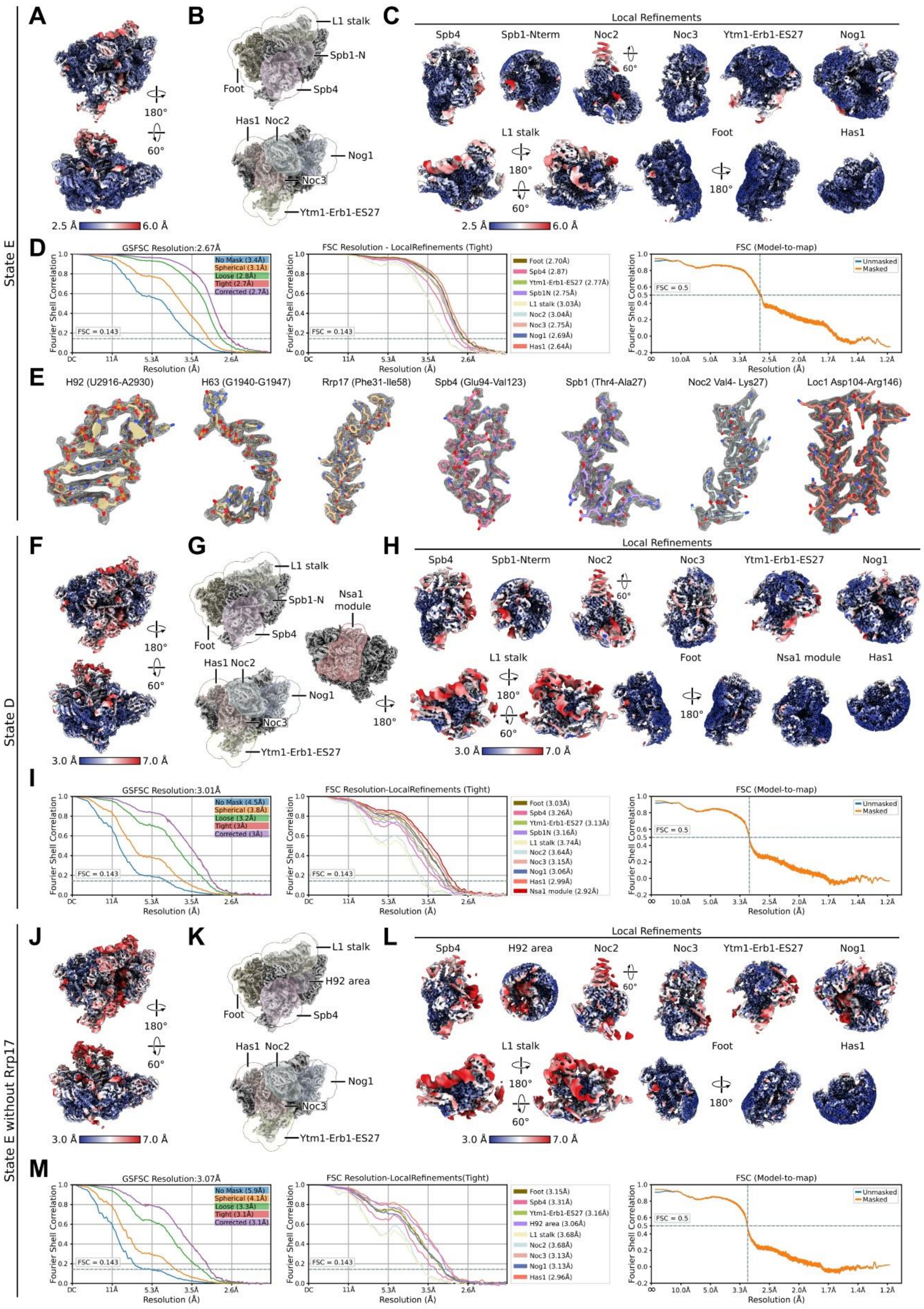

**Figure3–Figure Supplement 2. Local resolution distribution and FSC curves for state E, state D and state E lacking Rrp17.** Shown are data for state E (A-E), state D (F-I) and the state E without Rrp17 (J-M). (A, F, J) Consensus cryo-EM map of state E, state D and the state E lacking Rrp17 coloured according to local resolution. (B, G, K) The different consensus cryo-

EM maps and masks used for the local refinements. (C, H, L) Local refinements coloured according to local resolution. (D, I, M) Fourier shell correlation (FSC) of the consensus map (left), local refined maps (middle) and the model-to-map-FSC curves (right). (E) Segmented densities shown as mesh and corresponding models of individual parts of the nucleolar state E particle.

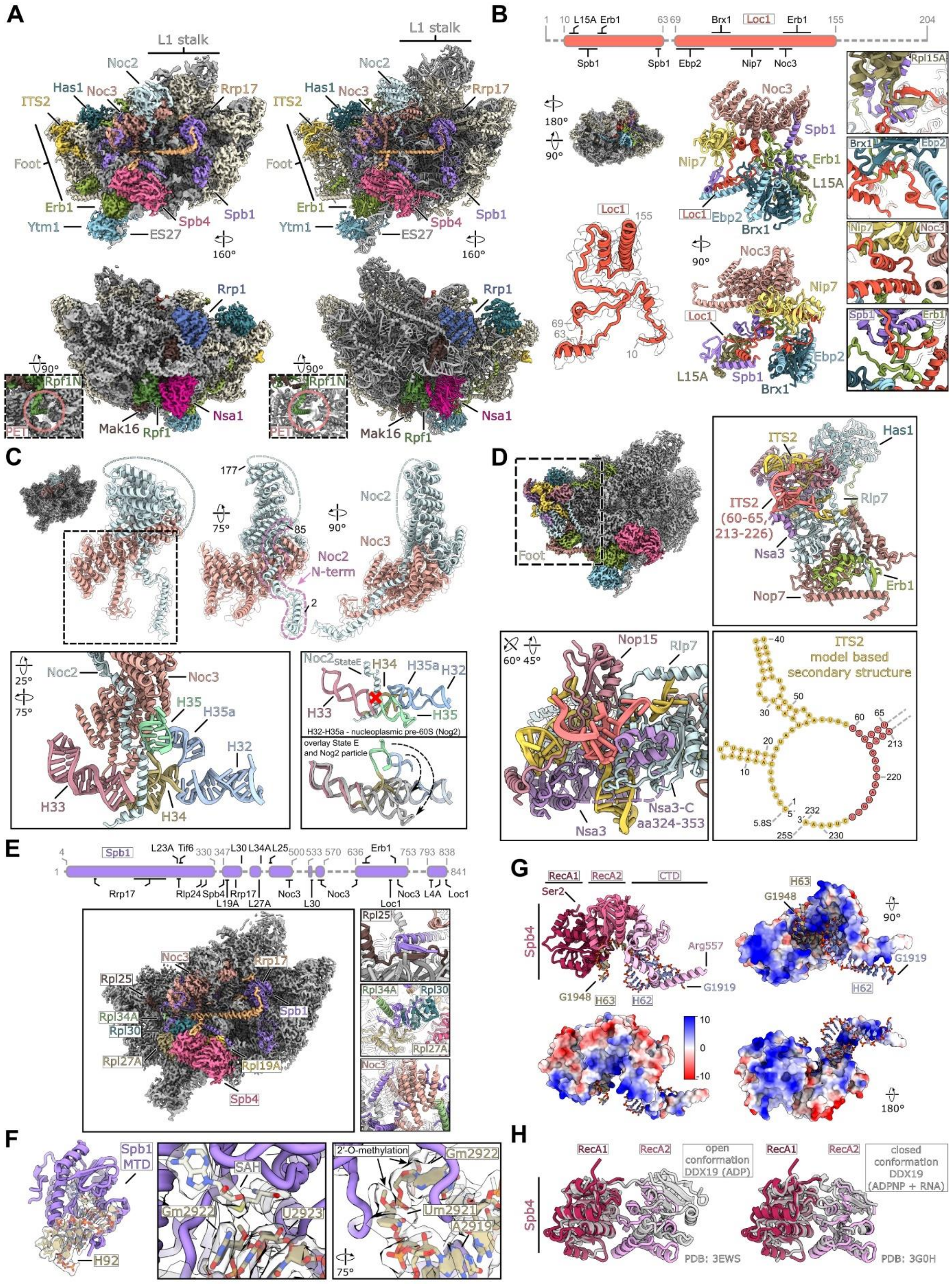


**Figure3–Figure Supplement 3. Structural details of nucleolar pre-60S states D/E.** (A) Cryo-EM density (left panels) and model (right panels) of the nucleolar pre-60S state D containing the Nsa1 module (Nsa1–Rpf1–Mak16–Rrp1). Close-up views show the N-terminus of Rpf1 entering the polypeptide exit tunnel (PET). (B) The Loc1 interaction network. Schematic

representation of the Loc1 interactions, unmodelled regions of Loc1 are shown as dashed lines (upper panel). Cryo-EM density of the state E class with the highlighted Loc1 interaction network and the molecular model and segmented density of Loc1 (left panels). Interaction network of the intrinsic Loc1 in two orientations. Only interacting regions and domains of the interacting factors are shown for simplicity (middle panels). Magnified regions of selected interactions between Loc1 and interacting proteins (right panels). (C) The Noc2–Noc3 complex shown in three orientations (upper panels). The Noc2 N-terminus (aa 2-85) is highlighted and the unmodelled middle part (aa 86-176) is indicated as dashed line. Interaction of the Noc2 N-terminus with rRNA H35, H32-H35 (nt 802-941) within state E is shown in different colours in the lower left panel. The lower right panels compare the orientation of H32-H35 in the nucleoplasmic Nog2 particle and the nucleolar state E. The sterical clash between the nucleoplasmic rRNA (Nog2 particle) and the Noc2 N-terminus (State E) is indicated showing that Noc2 release is required for H35 and H35a maturation. The H32-H35a rRNA of the Nog2 state (grey) is superimposed with the H32-H35a rRNA within state E (coloured). The rearrangements of H35 and H35a are indicated by arrows. (D) Cryo-EM map of the state E focussing on the ITS2-foot area. Magnifications of the foot-ITS2 region in different orientation are shown and the new modelled region of the ITS2 (nt 60-65 and nt 213-226) is indicated in red. Secondary structure of the ITS2 based on the state E model (lower right panel). (E) Interaction network of Spb1 with biogenesis factors and ribosomal proteins identified in this study. A schematic overview of Spb1 with its various interaction partners is shown. Unmodelled parts are shown as dashed line (upper panel). Spb1 is shown as cartoon representation and the state E subunit is shown as cryo-EM density. Interacting proteins identified in this study are coloured and labelled (lower left panel). Magnified regions showing selected interactions of Spb1 with Rpl25, Rpl30, Rpl34A and Noc3 are shown (lower right). (F) The Spb1-MTD interacting with the A loop H92 (left panel). Close-ups of SAH and the methylated target Gm2922 in two orientations. Surrounding nucleotides including Um2921 are labelled (middle and right panel). The segmented densities for SAH and H92 are shown and the 2′-O-methylations of Gm2922 and Um2921 are highlighted. (G) Model of Spb4 interacting with the immature rRNA H62 and H63 (upper left panel) and surface views of Spb4 coloured by its electrostatic potential and in complex with H62-H63 are shown in different orientations. (H) The RecA domains of Spb4 are in a closed conformation in state D/E. Comparison of the Spb4 helicase core model with superimposed models of DDX19 in an open conformation bound to ADP (PDB-3EWS) and in the closed conformation bound to ATP-analogue (ADPNP) and target RNA (3G0H). The DDX19 models were fitted to the RecA1 domain of Spb4. The CTD of Spb4, the N-terminus of DDX19 as well as nucleotides and target RNAs are omitted for clarity.

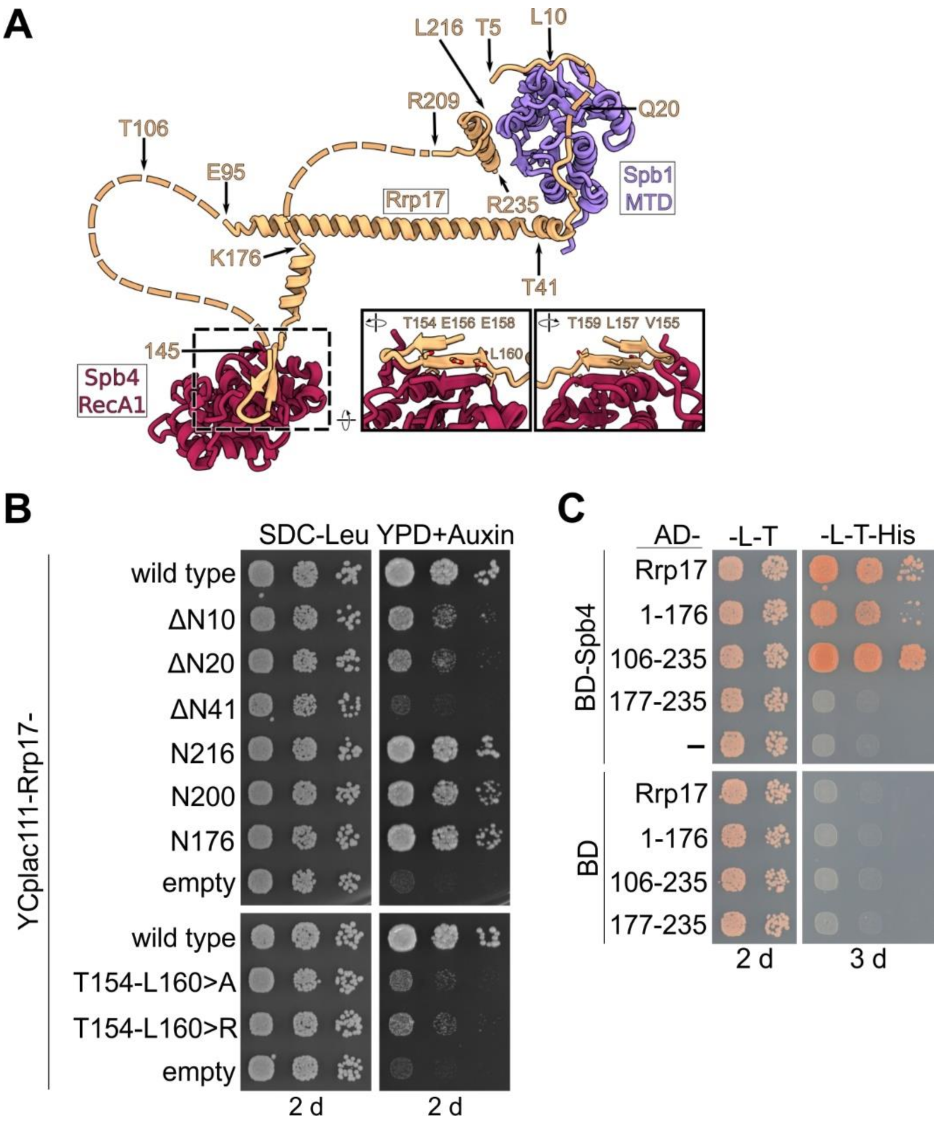


**Figure 3–Figure Supplement 4. Importance of the Spb4–Rrp17 β-sheet augmentation for cell growth and protein–protein interaction.** (A) Rrp17 in contact with the Spb1-MTD and Spb4-RecA1 domains in state E pre-60S subunits. Truncations and mutations of Rrp17 used for growth complementation and Y2H studies (B and C) are indicated. (B) An *RRP17*-HA-AID strain was transformed with plasmids harbouring wild-type *RRP17* or the indicated *rrp17* mutant alleles. Transformants were spotted in 10-fold serial dilutions on SDC-Leu plates or YPD plates containing 2 mM auxin (YPD+Auxin) and growth monitored after incubation for 2 days at 30 °C. (C) Spb4 fused to the Gal4 DNA-binding domain (BD) and indicated Rrp17 constructs fused to the Gal4 activation domain (AD) were transformed into the PJ69-4A yeast two-hybrid strain. Cells were spotted in ten-fold serial dilutions on SDC-Leu-Trp (-L-T) and SDC-Leu-Trp-His (-L-T-His; growth on this medium indicates a yeast two-hybrid interaction) plates and growth was monitored after incubation at 30 °C for 2 and 3 days, respectively.

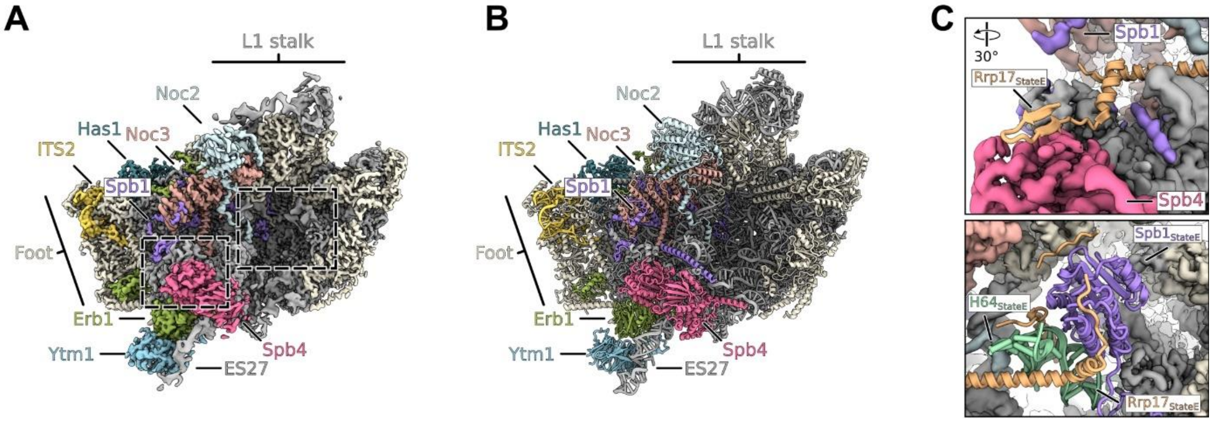


**Figure 3–Figure Supplement 5. Cryo-EM structure of the state preceding state E lacking Rrp17 and the Spb1-MTD.** (A-B) Coloured map (A) and model (B) of the state E like state lacking Rrp17 and the N-terminal MTD domain of Spb1. Relevant factors are coloured and labelled. (C) Coloured cryo-EM density of the state lacking Rrp17 focusing on the Spb4 RecA1 region (upper panel) and the region where the Spb1-MTD domain binds in state E (lower panel). The models of Rrp17, the Spb1-MTD and rRNA H64 of the state E were overlaid.

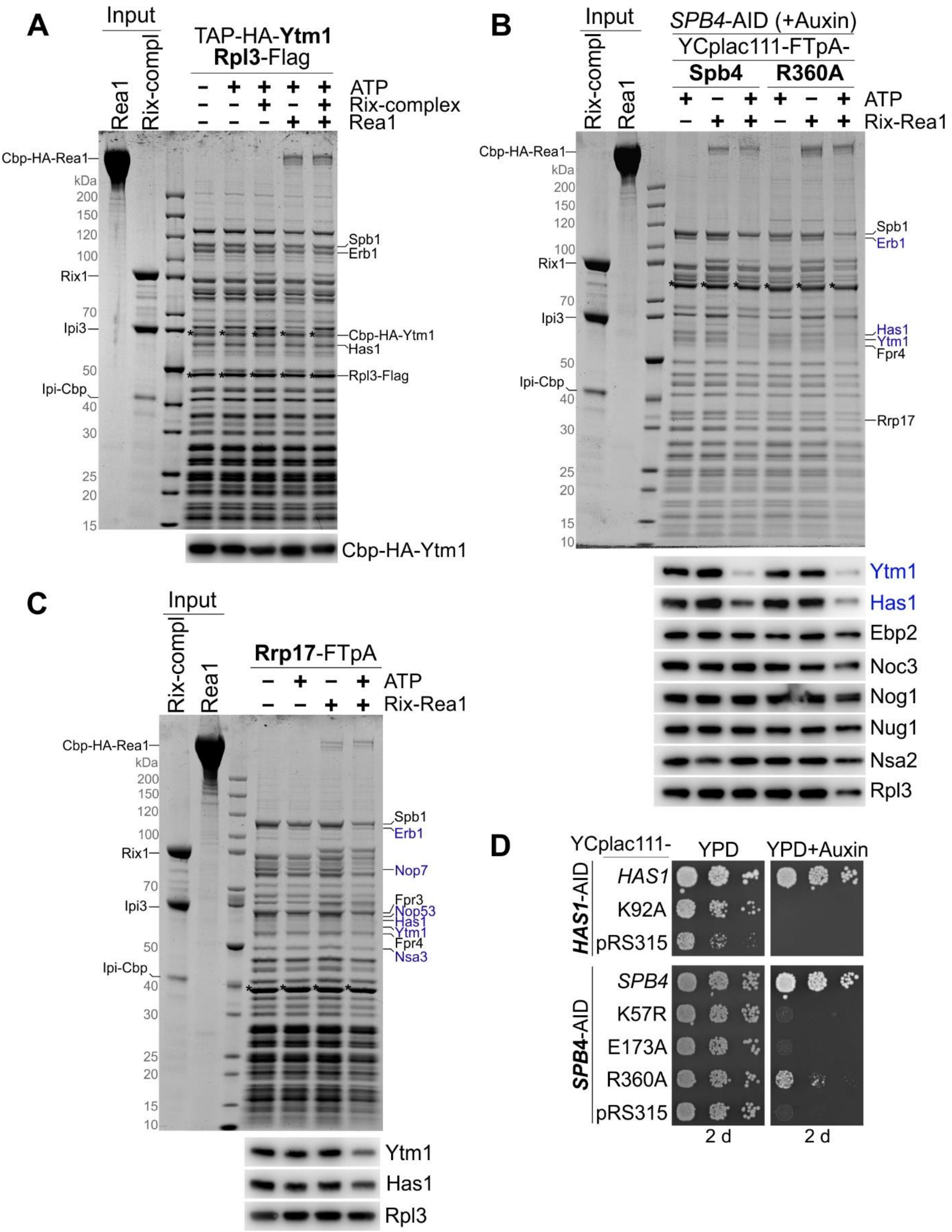


**Figure 4–Figure Supplement 1. *In vitro* maturation assays utilizing distinct substrate particles.** (A) Rea1-dependent remodelling cannot be induced on Ytm1-derived pre-60S particles. TEV-eluates of pre-60S particles isolated via TAP-HA-Ytm1 were as indicated incubated with purified Rea1, Rix1-complex ($Rix1_2$–$Ipi3_2$–Ipi1) and/or 3 mM ATP for 45 min at room temperature. Still bound material was re-isolated via Rpl3-Flag by incubation with anti-

Flag agarose beads and eluted with Flag peptide. Final eluates were analysed by SDS-PAGE and Coomassie staining (upper panel) or Western blotting using an anti-HA antibody to detect Ytm1 (lower panel). (B) Ytm1–Erb1 and Has1 are released from Spb4 R360A-derived mutant particles. Plasmid-borne FTpA tagged wild-type and R360A mutant Spb4 proteins were isolated from cells upon auxin-induced depletion of chromosomal *SPB4*-HA-AID. TEV eluates were incubated with Rea1, Rix1-complex and/or 3 mM ATP as described above and re-isolated via Spb4-Flag. Final eluates were analysed by SDS-PAGE and Coomassie staining (upper panel) or Western blotting with the indicated antibodies (lower panel). (C) TEV-eluates of pre-60S particles isolated via Rrp17-FTpA were incubated with Rea1, Rix1-complex and/or 3 mM ATP as described above and re-isolated via Rrp17-Flag. Final eluates were analysed by SDS-PAGE and Coomassie staining (upper panel) or Western blotting with the indicated antibodies (lower panel). Released proteins are indicated in blue. The bait proteins are marked with an asterisk. (D) A *HAS1*-HA-AID strain was transformed with plasmids harbouring wild-type *HAS1* or the *has1* K92A allele (upper panel) and a *SPB4*-HA-AID strain was transformed with plasmids harbouring indicated *spb4* alleles (lower panel). Transformants were spotted in 10-fold serial dilution on YPD plates with or without auxin and growth assessed after incubation at 30 °C for 2 days.

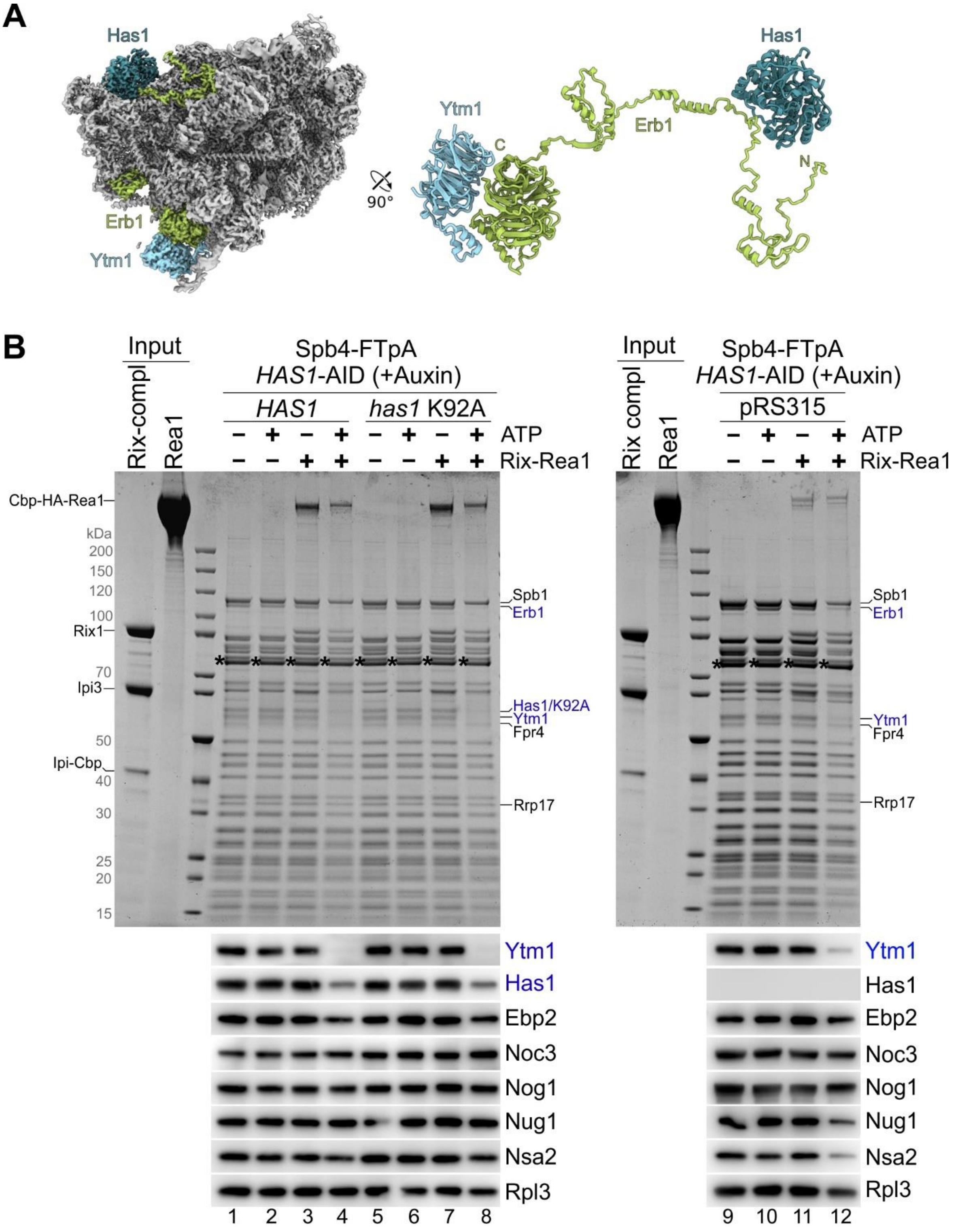


**Figure 4–Figure Supplement 2. Catalytically inactive Has1 K92A protein is dislodged from Spb4 particles upon Ytm1–Erb1 release.** (A) Cryo-EM reconstruction of the nucleolar state E pre-60S particle showing the contact between Has1 and the long unstructured Erb1 N-terminal tail. Ytm1 (light blue), Erb1 (green), and Has1 (blue) are displayed bound to the pre-60S particle (grey) as coloured cryo-EM map (left panel) or enlarged as cartoon representation (right panel). (B) Spb4-FTpA particles were isolated upon *HAS1*-HA-AID depletion from cells

expressing wild-type *HAS1* or a *has1* K92A Walker A mutant (left panel) or in absence of plasmid-borne *HAS1* (right panel). Spb4 TEV eluates were as indicated incubated with purified Rea1, Rix1-complex ($Rix1_2$–$Ipi3_2$–Ipi1) and/or 3 mM ATP for 45 min at room temperature. Still bound material was re-isolated via Spb4-Flag by incubation with anti-Flag agarose beads and eluted with Flag peptide. Final eluates were analysed by SDS-PAGE and Coomassie staining (upper panels) or Western blotting using the indicated antibodies (lower panels). The released proteins are indicated in blue.

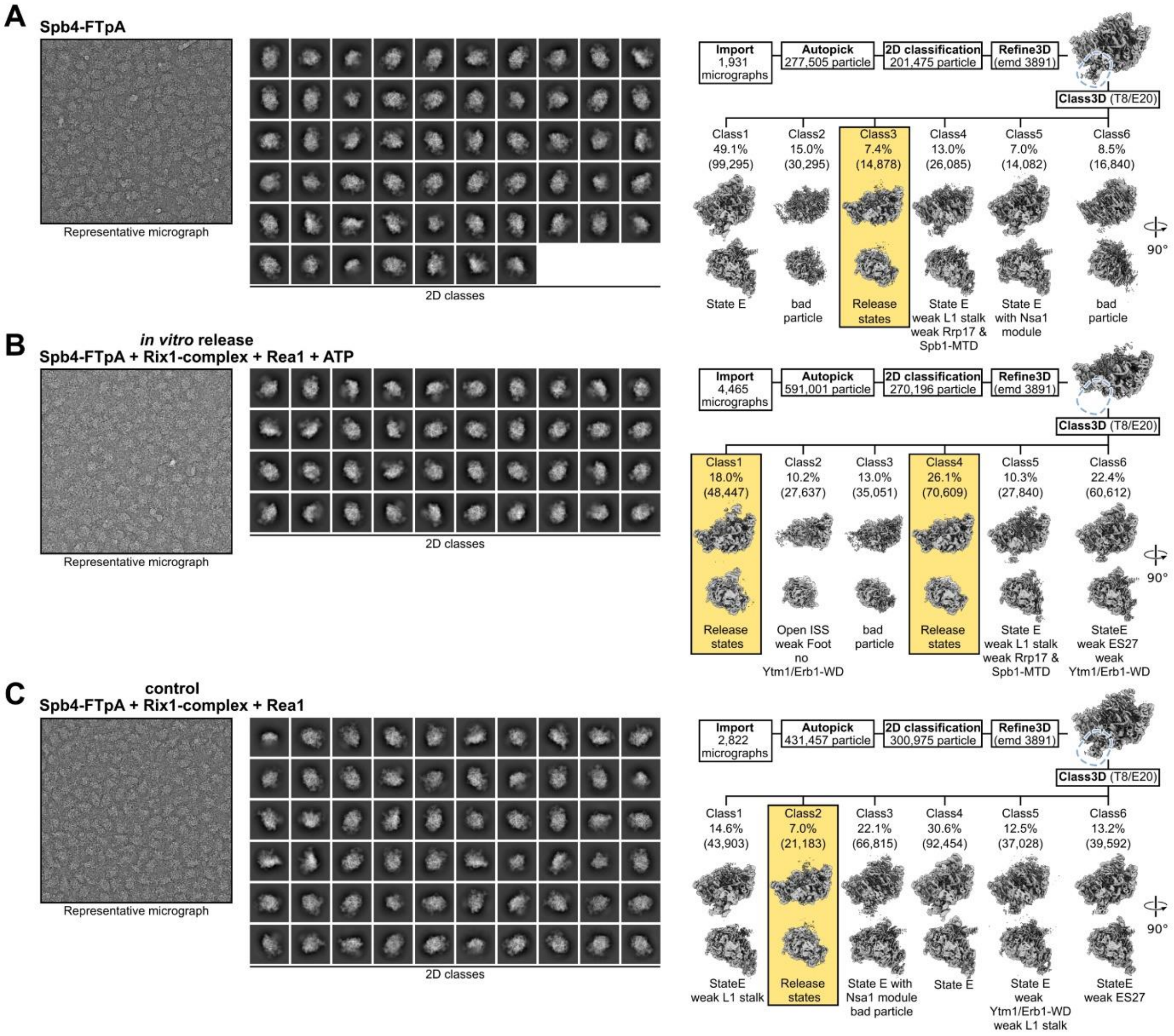


**Figure 4–Figure Supplement 3. Comparison of Spb4, *in vitro* release and control samples.** Processing schemes for the (A) Spb4-FTpA sample, (B) the *in vitro* release sample (Spb4-FTpA + Rix1 complex + Rea1 and ATP) and (C) the control sample (Spb4-FTpA + Rix1 complex + Rea1 but lacking ATP). The left and middle panels show representative micrographs and 2D class averages, respectively. The right panels show the initial processing schemes. After the initial refinement (Refine3D) density for the Ytm1-WD and Erb1-WD complex is clearly visible in the Spb4 and control samples but absent in the *in vitro* release sample (indicated with blue dashed lines). Subsequent 3D classification for each dataset is shown and the percentage and particle numbers (in brackets) are shown. The release state classes of each dataset are highlighted.

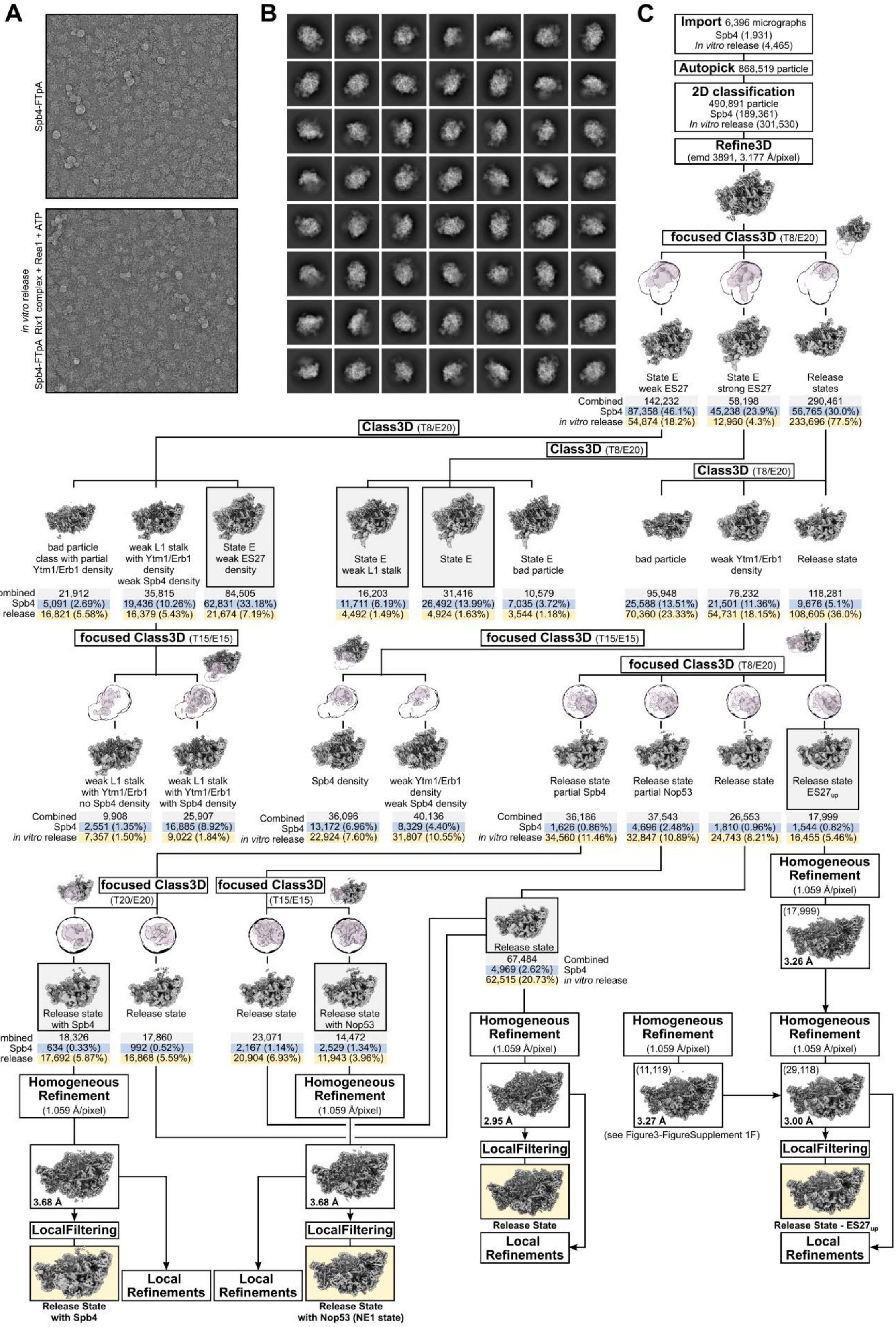


**Figure 4–Figure Supplement 4. Sorting scheme of the combined Spb4 and *in vitro* release datasets.** Representative micrographs of each dataset (A) and combined particle averages after 2D classification (B). (C) Sorting scheme of the combined dataset. Particle

numbers of the combined and the individual dataset of each class are shown. Final volumes are highlighted and shown with their respective resolution. In case of the release state ES27up volume, particles from an identical class (see Figure 3–Figure Supplement 1F) were added to increase the resolution.

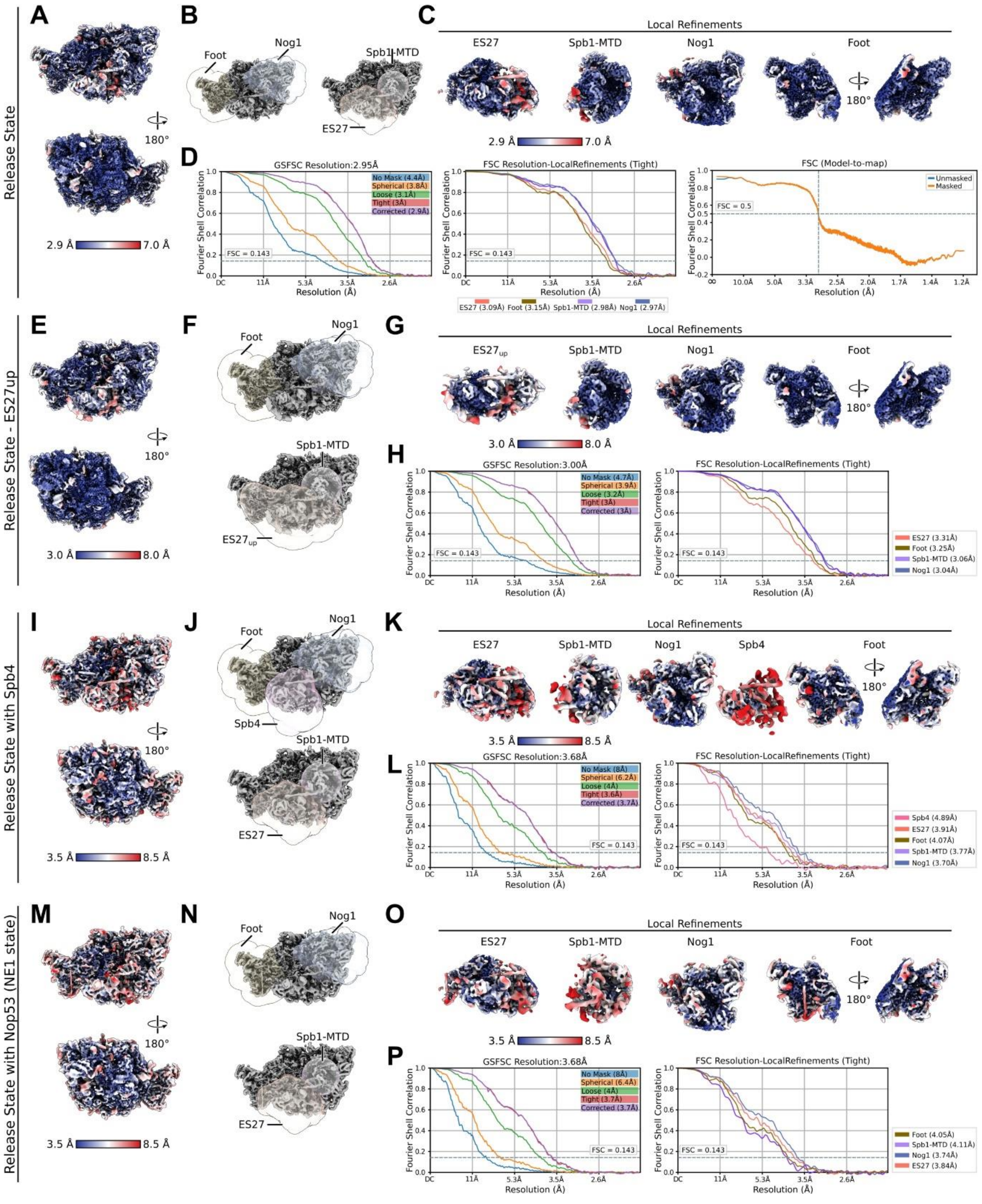

**Figure 4–Figure Supplement 5. Local resolution and FSC curves of the observed release states.** Data for the release state (A-D), the release state – ES27up (E-H), the release state with Spb4 (I-L) and the NE1 state containing Nop53 (M-P) are depicted. Consensus cryo-EM maps coloured according to the local resolution (A, E, I, M), the different consensus cryo-EM maps superimposed with masks used for the local refinements (B, F, J, N) and the different local refinements coloured according to the local resolution (C, G, K, O) are displayed. The Fourier shell correlation (FSC) curves for the consensus map (left) and the local refined maps (D, H, L, P) as well as the model-to-map-FSC curve for the release state (D) are shown.

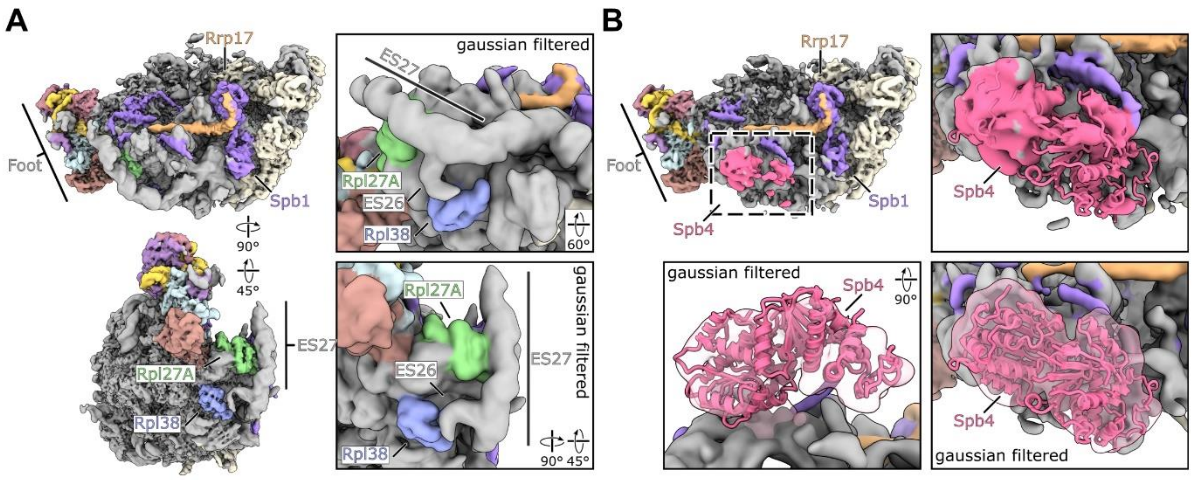


**Figure 4–Figure Supplement 6. Cryo-EM maps of the release state with ES27up and the release state containing Spb4.** (A) Coloured cryo-EM map of the release state with ES27 in an upward conformation shown in two orientations (left panels). Gaussian filtered map focussing on contact sites between Rpl27A-ES27 and Rpl38-ES26 (right panels). (B) Cryo-EM map overview of the Spb4 containing release state and close-up views focussing on the Spb4 density. Gaussian filtered maps are indicated. The model of the Spb4 helicase core (aa 2-398) was rigid body fitted into the density.

**Supplementary File 1. Yeast strains used in this study**

| Name | Genotype | Source |
|---|---|---|
| W303 | *ade2-1, his3-11,15, leu2-3,112, trp1-1, ura3-1, can1-100* | Thomas and Rothstein,1989 |
| *SPB4*-FTpA | W303 *MAT***a** *SPB4*-FTpA::nat*NT2* | this study |
| *RRP17*-FTpA | W303 *MAT***a** *RRP17*-FTpA::nat*NT2* | this study |
| *SPB4*-HA-AID | W303 *MAT*α *SPB4*-HA-AID:: HIS3MX6 P.*ADH1*-Os*TIR1*-9xmyc::*TRP1* | this study |
| *RRP17*-HA-AID | W303 *MAT*α *RRP17*-HA-AID:: HIS3MX6 P.*ADH1*-Os*TIR1*-9xmyc::*TRP1* | this study |
| *SPB4*-HA-AID *NOP7*-FTpA | W303 *MAT*α *SPB4*-HA-AID:: HIS3MX6 P.*ADH1*-Os*TIR1*-9xmyc::*TRP1* *NOP7*-FTpA::natNT2 | this study |
| *SPB4*-HA-AID *NUG1*-FTpA | W303 *MAT*α *SPB4*-HA-AID:: HIS3MX6 P.*ADH1*-Os*TIR1*-9xmyc::*TRP1* *NUG1*-FTpA::natNT2 | this study |
| *SPB4*-HA-AID *ARX1*-FTpA | W303 *MAT*α *SPB4*-HA-AID:: HIS3MX6 P.*ADH1*-Os*TIR1*-9xmyc::*TRP1* *ARX1*-FTpA::natNT2 | this study |
| *RRP17*-HA-AID *NOP7*-FTpA | W303 *MAT*α *RRP17*-HA-AID:: HIS3MX6 P.*ADH1*-Os*TIR1*-9xmyc::*TRP1* *NOP7*-FTpA::natNT2 | this study |
| *RRP17*-HA-AID *NUG1*-FTpA | W303 *MAT*α *RRP17*-HA-AID:: HIS3MX6 P.*ADH1*-Os*TIR1*-9xmyc::*TRP1* *NUG1*-FTpA::natNT2 | this study |
| *RRP17*-HA-AID *ARX1*-FTpA | W303 *MAT*α *RRP17*-HA-AID:: HIS3MX6 P.*ADH1*-Os*TIR1*-9xmyc::*TRP1* *ARX1*-FTpA::natNT2 | this study |
| TAP-HA-*YTM1* *RPL3*-Flag | W303 *MAT*α TAP-HA-*YTM1*::natNT2 *RPL3*-Flag::HIS3MX6 | this study |
| *HAS1*-HA-AID | W303 *MAT*α *HAS1*-HA-AID:: HIS3MX6 P.*ADH1*-Os*TIR1*-9xmyc::*TRP1* | this study |
| *HAS1*-HA-AID *SPB4*-FTpA | W303 *MAT*α *HAS1*-HA-AID:: HIS3MX6 P.*ADH1*-Os*TIR1*-9xmyc::*TRP1* *SPB4*-FTpA::natNT2 | this study |
| *SPB4* Shuffle | W303 *MAT***a** *spb4*::kanMX [Ycplac33-*SPB4*] | this study |
| PJ69-4A | *trp1-901, leu2-3,112, ura3-52, his3-200, gal4Δ, gal80Δ, LYS2::GAL1- HIS3, GAL2-ADE2, met2::GAL7-lacZ* | James *et al*. 1996 |

**Supplementary File 2. Plasmids used in this study**

| Name | Relevant Information | Source |
|---|---|---|
| YCplac111-TAP-Flag-*YTM1* | CEN, *LEU2,* P*YTM1, YTM1*, N-terminal TAP-Flag tag | Kater *et al*., 2017 |
| YCplac111-TAP-Flag-*ytm1*E80A | CEN, *LEU2,* P*YTM1, ytm1*E80A, N-terminal TAP-Flag tag | Kater *et al*., 2017 |
| YCplac111-*SPB4* | CEN, *LEU2,* P*SPB4, SPB4* | this study |
| YCplac111-*spb4*K57R | CEN, *LEU2,* P*SPB4, spb4*K57R | this study |
| YCplac111-*spb4*E173A | CEN, *LEU2,* P*SPB4, spb4*E173A | this study |
| YCplac111-*spb4*R360A | CEN, *LEU2,* P*SPB4, spb4*R360A | this study |
| YCplac111-*spb4*N563 | CEN, *LEU2,* P*SPB4, spb4*N563 (aa 1-563) | this study |
| YCplac111-*spb4*N531 | CEN, *LEU2,* P*SPB4, spb4*N531 (aa 1-531) | this study |
| YCplac111-*spb4*N470 | CEN, *LEU2,* P*SPB4, spb4*N470 (aa 1-470) | this study |
| YCplac111-*spb4*N405 | CEN, *LEU2,* P*SPB4, spb4*N405 (aa 1-405) | this study |
| YCplac111-*SPB4*-FTpA | CEN, *LEU2,* P*SPB4, SPB4*, C-terminal FTpA tag | this study |
| YCplac111-*spb4*K57R-FTpA | CEN, *LEU2,* P*SPB4, spb4*K57R, C-terminal FTpA tag | this study |
| YCplac111-*spb4*E173A-FTpA | CEN, *LEU2,* P*SPB4, spb4*E173A, C-terminal FTpA tag | this study |
| YCplac111-*spb4*R360A-FTpA | CEN, *LEU2,* P*SPB4, spb4*R360A, C-terminal FTpA tag | this study |
| YCplac111-*spb4*N563-FTpA | CEN, *LEU2,* P*SPB4, spb4*N563, C-terminal FTpA tag | this study |
| YCplac111-*spb4*N470-FTpA | CEN, *LEU2,* P*SPB4, spb4*N470, C-terminal FTpA tag | this study |
| YCplac111-*spb4*N405-FTpA | CEN, *LEU2,* P*SPB4, spb4*N405, C-terminal FTpA tag | this study |
| YCplac111-*RRP17* | CEN, *LEU2,* P*SPB4, RRP17* | this study |
| YCplac111-*rrp17*ΔN10 | CEN, *LEU2,* P*SPB4, rrp17*ΔN10 (aa 11-235) | this study |
| YCplac111-*rrp17*ΔN20 | CEN, *LEU2,* P*SPB4, rrp17*ΔN20 (aa 21-235) | this study |
| YCplac111-*rrp17*ΔN41 | CEN, *LEU2,* P*SPB4, rrp17*ΔN41 (aa 42-235) | this study |
| YCplac111-*rrp17*N216 | CEN, *LEU2,* P*SPB4, rrp17*N216 (aa 1-216) | this study |
| YCplac111-*rrp17*N200 | CEN, *LEU2,* P*SPB4, rrp17*N200 (aa 1-200) | this study |

| | | |
|---|---|---|
| YCplac111-*rrp17*N176 | CEN, *LEU2,* P*SPB4, rrp17*N176 (aa 1-176) | this study |
| YCplac111-*rrp17*T154-L160>A | CEN, *LEU2,* P*SPB4, rrp17*T154-L160>7xA | this study |
| YCplac111-*rrp17* T154-L160>R | CEN, *LEU2,* P*SPB4, rrp17*T154-L160>7xR | this study |
| YCplac111-*HAS1* | CEN, *LEU2,* P*HAS1, HAS1* | this study |
| YCplac111-*has1*K92A | CEN, *LEU2,* P*HAS1, has1*K92A | this study |
| YCplac111-P*GAL1-10*-TAP-HA-*REA1* | CEN, *LEU2*, P*GAL1-10*, N-terminal TAP-HA tag | this study |
| YEplac112-P*GAL1-10-RIX1*-TEV-pA | 2µ, *TRP1*, P*GAL1-10*, C-terminal TEV-pA tag | Barrio-Garcia *et al*., 2016 |
| YEplac181-P2-*IPI1*-Cbp-P*GAL1-10*-P1-*IPI3* | 2µ, *LEU2*, P*GAL1-10*, C-terminal Ipi1 Cbp tag | this study |
| pG4BDC22-*SPB4* | CEN, *TRP1*, P*ADH1*, C-terminal Gal4-BD | this study |
| pG4ADN111-*RRP17* | CEN, *LEU2*, P*ADH1*, N-terminal Gal4-AD | this study |
| pG4ADN111-*rrp17* (1-176) | CEN, *LEU2*, P*ADH1*, N-terminal Gal4-AD | this study |
| pG4ADN111-*rrp17* (106-235) | CEN, *LEU2*, P*ADH1*, N-terminal Gal4-AD | this study |
| pG4ADN111-*rrp17* (177-235) | CEN, *LEU2*, P*ADH1*, N-terminal Gal4-AD | this study |
